# Cell-surface *N*-linked glycans manipulation of K562 cells for augmented susceptibility to natural killer cell killing

**DOI:** 10.64898/2026.08.09.743798

**Authors:** Zhengyuan Huang, Qiqi Li, Alex T. H. Cocker, Hugh Brady, Mark R. Johnson

**Affiliations:** Department of Metabolism, Digestion and Reproduction, Imperial College London, SW10 9NH, London, United Kingdom; Department of Life Sciences, Imperial College London, SW7 2AZ, London, United Kingdom

**Keywords:** K562, *N*-glycosylation, sialylation, HLA-G, NK susceptibility

## Abstract

Glycosylation of proteins is arguably the most diverse post-translational modification that is altered in almost all cancer types, which has been demonstrated to play a crucial role in creating an immunosuppressive microenvironment that promotes immune tolerance and evasion. However, the biosynthesis of *N*-linked glycan is mediated by a series of enzymatic reactions catalysed by glycosyltransferases and glycosidases in a template-independent manner, hindering our understanding of specific structure-function relationships and roles of specific glycans on specific proteins. Here we use HLA class I-negative cell line K562, a known reference target for NK-mediated cytolysis, to establish a model investigating how cell surface glycan dynamics influence its susceptibility to cytolysis mediated by NK-92 cells. Treatment of K562 cells with kifunensine, swainsonine, 2F-peracetyl-fucose, or 3Fax-peracetyl Neu5Ac, inhibitors of *N*-linked glycan processing, resulted in drastic alterations in cell surface carbohydrate phenotype, as could be shown by flow cytometric analysis of the lecti*N*binding properties of the cells. Despite these clear changes in carbohydrate phenotype, only K562 cells treated with either kifunensine or 3Fax-peracetyl Neu5Ac exhibited higher susceptibility to the cytolysis medidated by NK-92 cells accompanied with an increased CD107a expression by NK-92 cells. Although K562 cells overexpressing gene *MGAT3* exhibited a decreased NK-susceptibility, we further found that this decrease was not exclusively determined by the overexpression of gene *MGAT3* product bisecting β1,4-GlcNAc, because the treatment of 3Fax-peracetyl Neu5Ac reversed the resistance of K562 cell against NK-92 cell in despite of expressing higher levels of bisecting β1,4-GlcNAc. Expressing HLA-G on cell surface as extravillous trophoblast did not change the NK-susceptibility of K562 cells, despite evidence that HLA-G molecules expressed by K562 cells can bind to inhibitory receptor ILT2 expressed on NK-92 cell surface. These findings suggest that the level of terminal sialylation, outweighing other components in *N*-linked glycan, determines the NK-susceptibility of K562 cell, offering a new strategy to weaken the resistance of cancer cells so that the immune system can maximise the elimination.

## INTRODUCTION

Natural killer (NK) cells, as a component of the innate immune system, possess the capability to directly target and eliminate malignant cells [1,2]. They constitute a crucial first line of antitumor defense and represent a promising platform for cancer immunotherapy because of their intrinsic cytotoxic activity and their capacity to identify tumor cells independently of antigen presentation [3]. Tumor cells can evade NK cell activity, thereby limiting the antitumor immunity [4,5]. Dysregulated expression levels of glycosylatio*N*related enzyme have been reported to be associated with tumor development, progression, invasion, and metastasis [6–8]. Understanding the link between cell-surface glycan dynamics of malignant cell and its susceptibility to NK-mediated cytolysis (NK-susceptibility) has been alluring since Yoshimura et al found that human immortalized myelogenous leukemia cell line K562, which is served as a positive control for NK cell killing activity lacking major histocompatibility complexes required to inhibit NK activity [9], become resistant to the NK-mediated cytolysis through overexpressing *N*-glycosylation related gene *MGAT3*, along with an increase of *N*-linked glycan with bisecting β1,4-*N*-acetyl-glucosamine (GlcNAc, Fig. S1A) and decreases in terminal sialylation on cell surface [10]. The introduction of bisecting β1,4-GlcNAc has been reported to suppress further processing and elongation of *N*-linked glycans, precluding the formation of β1,6-GlcNAc branching catalysed by GnT-V (encoded by gene *MGAT5*), since GnT-V is unable to use the *N*-linked glycan with bisecting β1,4-GlcNAc as a substrate [11,12]. The ability of bisecting β1,4-GlcNAc to hinder terminal modifications of several terminal epitopes (e.g., fucose, sialic acid, and human natural killer-1) of bi-antennary *N*-linked glycans has also been reported in *Mgat3*-deficient mice [13]. Furthermore, based on a *MGAT3*-overexpressing human breast cancer cell line, the bisecting β1,4-GlcNAc is found to specifically inhibit the α2,3-rather than α2,6-sialylation [14]. The final *N*-linked glycan structures present on glycoproteins are produced through coordinated and competing activities of multiple glycosyltransferases in the Golgi apparatus, together with the existance of sugar likes bisecting β1,4-GlcNAc exhibiting suppressive effects in the biosynthesis of *N*-linked glycan, making it difficult to understand the mechanism that how glycosylation influences the NK-susceptibility of tumor cells.

While research on immune checkpoints has historically focused on protein factors [15], recent studies have highlighted glycosylated antigens as important drivers of cancer-associated immune suppression. Using CRISPR screens in multiple myeloma cell line LP-1, Dufva et al identified the protein fucosylation genes *FUT8*, *GMDS*, and *SLC35C1*, as well as the cell-surface protein gene *MUC1*, as key drivers of resistance to NK-mediated cytolysis [4]. Consistent with these findings, the C-terminal transmembrane subunit of muci*N*1 (O-glycosylated proteins encoded by gene *MUC1*) has also been shown to suppress NK cell-mediated cytotoxicity through repression of MICA/B in muci*N*expressing solid tumors [16]. Additional evidence upholding a role of glycosylation in NK cell-tumor interactions includes glycosylation regulator signal peptide peptidase-like 3, heavily glycosylated mucins, Siglec-7 ligands SPN (CD43), and SELPLG (PSGL-1) [17,18]. Notably, deletion of the gene encoding signal peptide peptidase-like 3 in the NK-sensitive B-cell lymphoblastoma cell line 721.221 resulted in enrichment of *N*-linked glycans with extended poly-*N*-acetyl-lactosamine (LacNAc) structures, thereby conferring increased resistance to NK-mediated cytolysis [19]. In conclusion, the remodeling of cell-surface glycans presumably changes the NK-susceptibility of tumor cell through affecting activities of glycoprotein involved in NK receptor binding.

Extravillous trophoblasts (EVTs), unlike K562 cells, display low antigenicity and do not induce a typical rejection response in immunocompetent hosts despite expressing rejection antigens and interacting extensively with decidual immune cells [20]. *N*-glycomic analyses of primary trophoblasts reveal distinct *N*glycomic profiles between extravillous and villous subpopulations [21]. Mouse knockout and transgenic studies indicate that trophoblast antigens bearing sialylated *N*-linked glycans suppress B-cell responses and promote fetomaternal tolerance, whereas their absence causes embryonic lethality due to strong complement activation against trophoblasts [22,23]. EVTs also exhibit a unique class I human leukocyte antigen (HLA-I) profile associated with fetomaternal tolerance, characterized by co-expression of classical HLA-C and no*N*classical HLA-E, -F, and -G [24,25], which is a defining feature of EVT differentiation [26,27]. These HLA molecules share a conserved *N*-glycosylation site in the heavy chain, which has been proved to be important for proper folding of HLA-Cw1 in the endoplasmic reticulum [28–30]. Using classical ^51^Cr cytotoxicity assay and peripheral blood NK cells present in PBMC from several healthy donors as effectors, Tronik-Le Roux et al demonstrated that K562 cell expressing HLA-G1 exhibited decreased NK-susceptibility compared to wildtype K562 cell [31]. Another study co-incubated decidual CD56^+^ NK cells with 721.221 cells transfected with HLA-G monomer or with the HLA-G homodimer and found that decidual NK cell produced cytokines conrtibuting to establish an immune-tolerant and pregnancy-supportive environment rather than mounting a cytotoxic immune attack [32].

Supposing that the NK-susceptibility of K562 cell can be modified through remodeling of cell-surface glycans, if there exist patterns that can fine tune its NK-susceptibility remain to be elucidated. In this regard, we here estabilsihed a simplified model using K562 cells with remodeling of cell-surface glycans to perform studies in relationship bettwen cell surface glycan structure and NK-susceptibility. The results revealed unique and common features of cell-surface glycan dynamics.

## MATERIALS AND METHODS

### Antibodies and reagents

Anti-HLA-G (clone 4H84) conjuageted with either Alexa Fluor™ 488 or horseradish peroxidase (HRP) was purchased from Santa Cruz Biotechnology, allophycocyanin (APC) or fluorescein isothiocyanate (FITC) anti-HLA-G antibody (clone MEM-G/9) from Abcam, anti-HLA-G antibody (clone G233) from Invitrogen, and HRP-conjugated secondary antibody against rabbit or mouse from Cell Signaling Technology. Brilliant Violet 650™ anti-human CD56 (NCAM) antibody, Brilliant Violet 421™ anti-human CD107a (LAMP-1) antibody, Direct-Blot™ HRP anti-β-actin or -GAPDH antibody, and streptavidin conjugated with either Brilliant Violet 421™ or phycoerythrin (PE) were all from BioLegend. Biotinylated lectins (Table S1) *Concanavalin* A (ConA), *Lycopersicon esculentum* Lectin (LEL), *Lens Culinaris* agglutinin (LCA), *Phaseolus vulgaris* erythroagglutinin (PHA-E), *Phaseolus Vulgaris* Leucoagglutinin (PHA-L), *Sambucus nigra* agglutinin (SNA) were all purchased from Vector Laboratories. All chemicals were purchased from Sigma-Aldrich (MA, USA) and media from Thermo Fisher Scientific (MA, USA), unless otherwise indicated.

### Cell Culture and glycomic remodeling

All cell lines were purchased from the American Type Culture Collection (VA, USA) and maintained at 37°C and 5% CO_2_ in a humid incubator. Floating cells were passaged when density recahed at 1×10^6^ cells/ml. Cell passages between 10-30 were used in this study. All media for cell culture were purchased from Gibco unless otherwise indicated.The human leukemia cell line K562 (CCL-243) and transgenic K562 cell overexpressing gene *MGAT3* conjugated to red fluorescent protein DsRed as a reporter (M3-K562), which was generated by the transfection with lentivirus vectors and gifted by Dr. Heather Ang (Departments of Life Sciences, Imperial College London, UK), were both cultured in RPMI 1640 medium supplemented with 10% (v/v) fetal bovine serum (FBS), 100 units/mL of penicillin, and 100 μg/mL of streptomycin (referring to “R10” hereafter). To achieve the predominance of either oligomannose or or hybrid *N*-linked glycans on cellular glycoproteins, K562 cells were cultured in the presence mannosidase inhibitors kifunensine (KIF) or swansonine (SWA) for 3 days, while the 3-day treatment of fucose analogue2F-peracetyl-fucose (2FF) and Neu5Ac analogue 3F_AX_-peracetyl-Neu5Ac (3FN) were respectively used to reduce the presence of sialylated and fucosylated *N*-linked glycans on cellular glycoproteins (Fig. S1A and Table S2) [18,33,34]. Alterations in cell surface carbohydrate phenotype were indcated by flow cytometric analysis of the lecti*N*binding properties of the cells. No*N*adherent immortal NK cell line NK-92 (CRL-2407), used as effector in cytotoxicity assays, was incubated in Minimum Essential Medium Eagle (Alpha Modification, M-0200, Sigma-Aldrich) supplemented with 0.2 mM inositol, 0.02 mM folic acid, 200 units/mL recombinant human IL-2, 12.5% (v/v) horse serum, 12.5% (v/v) FBS, 0.1 mM 2-mercaptoethanol (replenished daily), 100 units/mL of penicillin, and 100 μg/mL of streptomycin.

### First-trimester chorionic villi collection and protein extraction

First-trimester placental samples (6-13 weeks) were collected from women undergoing surgical termination at Chelsea and Westminster Hospital and West Middlesex University Hospital, with ethical approval (REC reference: 11/LO/0971). Samples were processed within 4 h of collection. Placental tissue was rinsed in ice-cold Dulbecco’s Phosphate-Buffered Saline (DPBS), dissected, and cleared of blood clots. Tissue was homogenized in RIPA buffuer using Precellys™ homogenizer (Bertin Technologies SAS, Montigny-le-Bretonneux, France), and then centrifuged at 10,000 × g for 10 min at 4°C. The supernatant was collected, mixed with Laemmli sample buffer containing 50 mM dithiothreitol, and heated at 96°C for 6 minutes.

### Proliferation assay

The cell proliferation during inhibition in glycosylation was studied using the CellTrace™ CFSE (carboxyfluorescein succinimidyl ester) Cell Proliferation Kit (Invitrogen) according to the manufacturer’s instruction. Approximately 1.5×10^5^ K562 cells were suspended in 1 µM CFSE staining solution, incubated at 37 for 20 minutes, rinsed with complete culture medium, and finally resuspended in fresh, pre-warmed complete culture medium in the presence or absence of inhibitors. For each condition, equal number of cell was seeded into three wells, allowing the CFSE intensity to be analysed by flow cytometry at 24, 48 and 72 hours after seeding.

### Glycomic profiling using lectin staining

Cells were washed with DPBS and resuspended in 100 μL of FACS staining buffer (DPBS supplemented with 1% FBS and 2 mM EDTA, pH 7.2). Cells were incubated with 10 μg/mL of biotinylated lectins (Vector Laboratories, CA, USA, Table S2) at 4°C for 15 minutes. Afterwards cells were washed with FACS staining buffer and combined with 10% (v/v) Precision Count Beads™ (BioLegend, CA, USA). Cells were analyzed by flow cytometry using a BD LSRFortessa™ cell analyzer (BD Biosciences, NJ, USA), and data analysis was performed using FlowJo™ software (Tree Star, CA, USA).

### Plasmid transfection in K562 cells using electroporation

K562 cells were transfected to express wildtype HLA-G (HLA-G^wildtype^), or HLA-G lacking its single *N*glycosylation site, which is created by changing the codon AAC for Asn110 (N110) to the codon CAG for Gln (Q) by referring to previous study [35], turning the *N*-glycosylation consensus YNQ into YQQ and is termed HLA-G^N110Q^ hereafter (Fig. S2A). K562 cells in logarithmic growth phase were harvested and resuspended with DPBS to give a density of approximately 5×10^4^ cells/µl, and 30 µl of this cell suspension was combined with 24 µl of gene expression vectors (0.5 µg/µl of each plasmid purchased from R&D systems) and mixed well with 54 µl of 2 × electroporation buffer from the Cell Line Nucleofector® Kit V (Lonza). This mixture was transferred into an electroporation cuvette, which was further placed into an Nucleofector^®^ 2b device (Lonza) to commence the electroporation. Afterwards, the cuvette was returned to the biosafety cabinet, where 500 µl of R-10 was added to the cuvette and mixed well using micro tip Pasteur pipettes. Afterwards, the entire content was transferred into a 25 cm^2^ culture flask containing 5.4 ml R-10 and incubated for 48 hours. The non-glycosylated HLA-G mutant was characterized by flow cytometry, immunoprecipitation, and Western blot upon transfection in K562. Different anti-HLA-G mAbs (Fig. S2B) were used to study the localization and expression pattern of HLA-G isoform as previous described but with modifications [36–38].

### SDS-PAGE and Western blotting

All reagents and equipment used for electrophoresis and Western blotting were obtained from Bio-Rad unless otherwise specified. Cells were lysed for 30 minutes in ice-cold RIPA buffer supplemented with 2 mM phenylmethylsulfonyl fluoride and 1 mM sodium orthovanadate (BioLegend). Protein concentrations of cell lysates from K562 cell transfectants were determined and normalized using the bicinchoninic acid assay (Pierce™ 660 nm Protein Assay Reagent and Pierce™ bovine serum albumin standard pre-diluted set, Thermo Fisher). Cell lysates were mixed with Laemmli sample buffer containing 50 mM dithiothreitol and heated at 96°C for 6 minutes. After cooling to room temperature, equal amounts of denatured proteins were separated by SDS-PAGE under reducing conditions and subsequently transferred onto PVDF membranes. Membranes were incubated with primary antibodies overnight and followed by incubation with HRP-conjugated secondary antibodies. Immunoreactive bands were visualized using Clarity Western ECL Substrate and the iBright™ FL1500 imaging system (Thermo Fisher). Data acquisition and analysis of immunoblots were performed using iBright™ Analysis Software (Thermo Fisher).

### Immunoprecipitation of HLA-G

All reagents and equipment used for immunoprecipitation were obtained from Invitrogen (MA, USA) unless otherwise specified. Dynabeads® Protein A/G were first coupled with anti-HLA-G antibody (clone G233) and incubated overnight at 4 °C with whole cell lysates of K562 cell transfectants on a tube rotator. HLA-G molecules were enriched by pelleting Dynabeads® Protein A/G complexes on a DynaMag™-2 magnet. The post-immunoprecipitation supernatant was collected and stored at −20 °C for further analysis, while the bead pellet was rinsed with DPBS for three times, resuspended in 80 μL of Laemmli sample buffer (Bio-Rad) supplemented with 50 mM dithiothreitol, heated at 96 °C for 6 minutes, and allowed to cool to room temperature before being placed back on the DynaMag™-2 magnet. After the beads formed a compact pellet, the supernatant containing HLA-G molecules was collected and stored at −20 °C for subsequent use.

### Flow cytometry-based cytotoxicity assay

Target cells were harvested and subjected to CFSE staining as describerd above. Both CFSE-stained target cells and NK-92 cells were resuspended in R-10 supplemented with 200 units/mL IL-2 to reach a density of 1.2-1.5 × 10^6^ cells/mL. 250 μL of each cell suspension was mixed to reach an effector-to-target ratio at 5:1 and incubated in a 24-well plate for 10 hours. In parallel, target cells were also cultured under the same condition in the absence of effector to determine their spontaneous death rate. At the end of co-culture, cells were harvested, washed, resuspended in FACS staining buffer, and then successively stained with LIVE/DEAD™ Fixable Near-IR Stain (L/D, dilution 1 in 1000), BV650-conjugated anti-CD56 mAb (dilution 1 in 100, BioLegend), and eventually subjected to viability assessment by flow cytometry. Debris and doublets were first excluded, and then target cells were identified by gating on CFSE^+^ CD56^−^. The L/D^+^ population was gated out, of which the percentage to the total CFSE^+^ CD56^−^ population represents the absolute death rate achieved for each condition. The cytotoxicity of NK-92 cell against target cell was evaluated as previously reported [39]:

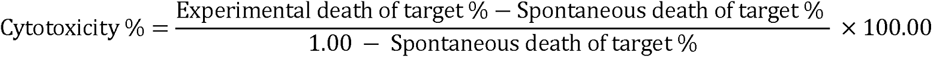

### CD107a degranulation assay

Degranulation evaluation of NK-92 cell was performed as previously reported [40]. K562 cells with or without glycomic remodeling were harvested, rinsed, resuspended in R-10 supplemented with 200 units/mL of IL-2, and co-incubated with NK-92 cells in a 5:1 effector-to-target ratio in 24-well plate, with a final volume of 500 μL per well. Preparing another two wells with NK-92 cell only for setting up positive and negative control (Fig. S1C, D). For positive control, 50 ng/mL phorbol 12-myristate 13-acetate (PMA) and 1 µg/mL ionomycin (Invitrogen) were additionally added. Subsequently, 1 µg of anti-human CD107a antibody was added into each well, and cells were incubated for 4 hours. After 1 hour of incubation, 2 μM of eBioscience™ monensin solution (Invitrogen) was added into each well and mixed by gentle pipetting. At the end of the 4-hour incubation, cells were harvested and subjected to flow cytometric analysis.

### Quantification and Statistical Analysis

Analysis was performed with Prism version 9.0.0 (GraphPad Software, Boston, MA). All data were tested for normality using a Kolmogorov-Smirnoff test or a D’Agostino & Pearson normality test depending on the group size. For normally distributed data, either paired or unpaired two-tailed Student’s t-test was used to perform comparison between two groups, while ANOVA followed by a Dunnett’s or Bonferroni’s post-hoc test for three groups or more. As for no*N*normally distributed data, either Wilcoxon signed-rank test or Man*N*Whitney U test was used to perform comparison between two paired or unpaired groups, while Friedman’s test with a Dunn’s multiple comparisons post-hoc test for three groups or more. Data are shown as mean ± SD based on at least three independent experiments. A *p*-value less than 0.05 was statistically significant (\**p* < 0.05), \*\**p* < 0.01, \*\*\**p* < 0.001, and \*\*\*\**p* < 0.0001.

## RESULTS

### The predominance of oligomannose *N*-linked glycan or de-sialylation results in an increased NK-susceptibility of K562 cells

The level of terminal α-mannose on K562 cell surface was enhanced by 4- to 6-fold after either KIF or SWA treatment, with concomitant absences in both bisecting β1,4-GlcNAc and β1,6-GlcNAc branches (Fig. 1A). Although both KIF- and SWA-treated K562 cells exhibited a remodelling of cell surface *N*-linked glycan, only the former one exhibited an increased NK-susceptibility (Fig. 1B). We also monitored the presence of activation marker CD107a on NK-92 cells during the co-culture with K562 cells since the trafficking of CD107a to the NK cell surface correlates with degranulation and the release of cytotoxic molecules such as perforin [41]. An increased expression of CD107a on NK-92 cell during co-incubation with KIF-treated K562 cell was observed (Fig. 1E). However, differences were found in α2,6-sialylation and core α1,6-fucosylation between these two inhibitor-treated samples, which were both markedly decreased in KIF-treated K562 cell when compared to that in control, while SWA-treated samples remained unaffected or showed the opposite (Fig. 1A). We therefore questioned if the level of sialylation or fucosylation is playing a more decisive role in determining the the NK-susceptibility of K562 cells compared to other glycan structures, especially the vital role of sialylation in regulating immunosuppression of immune cell has been abundantly demonstrated [10,18,42,43]. Although the 3FN treatment also induced increases in both NK-susceptibility of K562s cells and CD107a expression by NK-92 cell as KIF treatment (Fig. 1D and 1E), it only induced a significant reduction in SNA binding accompanied with increases of both bisecting β1,4-GlcNAc and β1,6-GlcNAc (Fig. 1C). As for the the 2FF treatment, it induced a significant reduction in core α1,6-fucosylation but increased the level of bisecting β1,4-GlcNAc branch, which did not affect either NK-susceptibility of K562 cells or CD107a expression by NK-92 cells (Fig. 1C-E). Notbaly, the proliferation of K562 cells was unaffected by the inhibition of glycosylation over three-day treatment, indicated by no significant difference in CFSE intensity and cell death between vehicle control and either inhibitor-treated sample (Fig. S1B).

**Figure 1.**
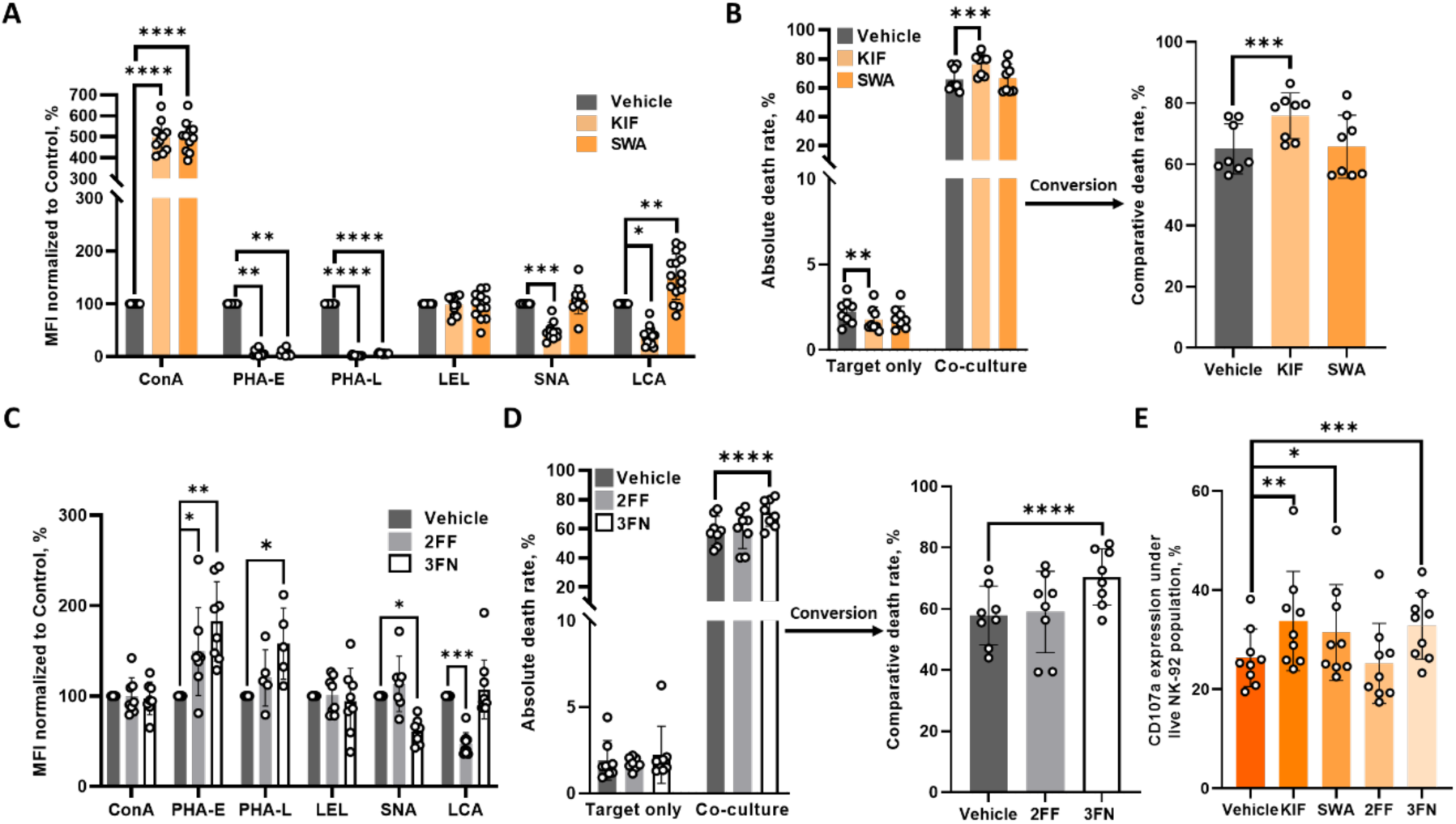
The NK-susceptibility of K562 cell can be augmented through the remodeling of cell-surface glycan. Flow cytometry of K562 cells using staining of representative lectins to compare the glycomic profile of cell surface between **(A)** vehicle control and KIF- or SWA-treated samples (N = 10-15), and between **(C)** vehicle control and 2FF- or 3FN-treated samples (N = 5-10). The MFI values of inhibitor-treated K562 cells were normalized to those of vehicle control. Same as below. Cytotoxicity assays of NK-92 cell co-incubated with K562 cell pretreated with **(B)** either KIF or SWA (N = 8), and with **(D)** either 2FF or 3FN (N = 8). The comparative death rates of vehicle and inhibitor-treated K562 cell after co-culture with five-fold of NK-92 cell were determined based on the experimental (“Co-culture”) and spontaneous (“Target only”) death rates of K562 cells. Same as below. **(E)** The CD107a expression of NK-92 cells after the co-culture with K562 cells possessing different profiles of cell-surface glycan. Paired Student’s t-test (*) was used to compare the means between vehicle control and either inhibitor-treated K562 cells.

### The low NK-susceptibility of M3-K562 cells can be reversed by the temporary remodeling of cell-surface glycan

Considering the effect of inhibitor on glycan processing can be temporary or reversible, we therefore sought for a remodeling of cell-surface glycan on K562 cell based on genetic modification. M3-K562 (transgeic K562 cell line which is overexpressing gene *MGAT3*) cell was exploited in this study to further compare the influences in NK-susceptibility of K562 cells between inherent and temporaty remodeling of cell-surface glycan. In agreement with previous studies by Yoshimura et al [9], M3-K562 cell exhibited a decresed NK-susceptibility compared to wildtype K562 cell (Fig. 2A) in flow cytometry-based cytotoxicity assays, but did not change the CD107a expression by NK-92 cells (Fig. 2B). Further comparison in glycomic profile between wildtype K562 and M3-K562 cells showed that those *N*-linked glycans present on M3-K562 cell surface contain higher levels of both bisecting β1,4-GlcNAc and β1,6-GlcNAc branches, but lower level of α2,6-sialylation (Fig. 2C). However, although the 3FN treatment can further increased the levels of both bisecting β1,4-GlcNAc and β1,6-GlcNAc branches while resulted in a lower level of α2,6-sialylation on M3-K562 cell surface (Fig. 2D), M3-K562 cells with 3FN treatment still exhibited an increased NK-susceptibility (Fig. 2E) and CD107a expression (Fig. 2F).

**Figure 2.**
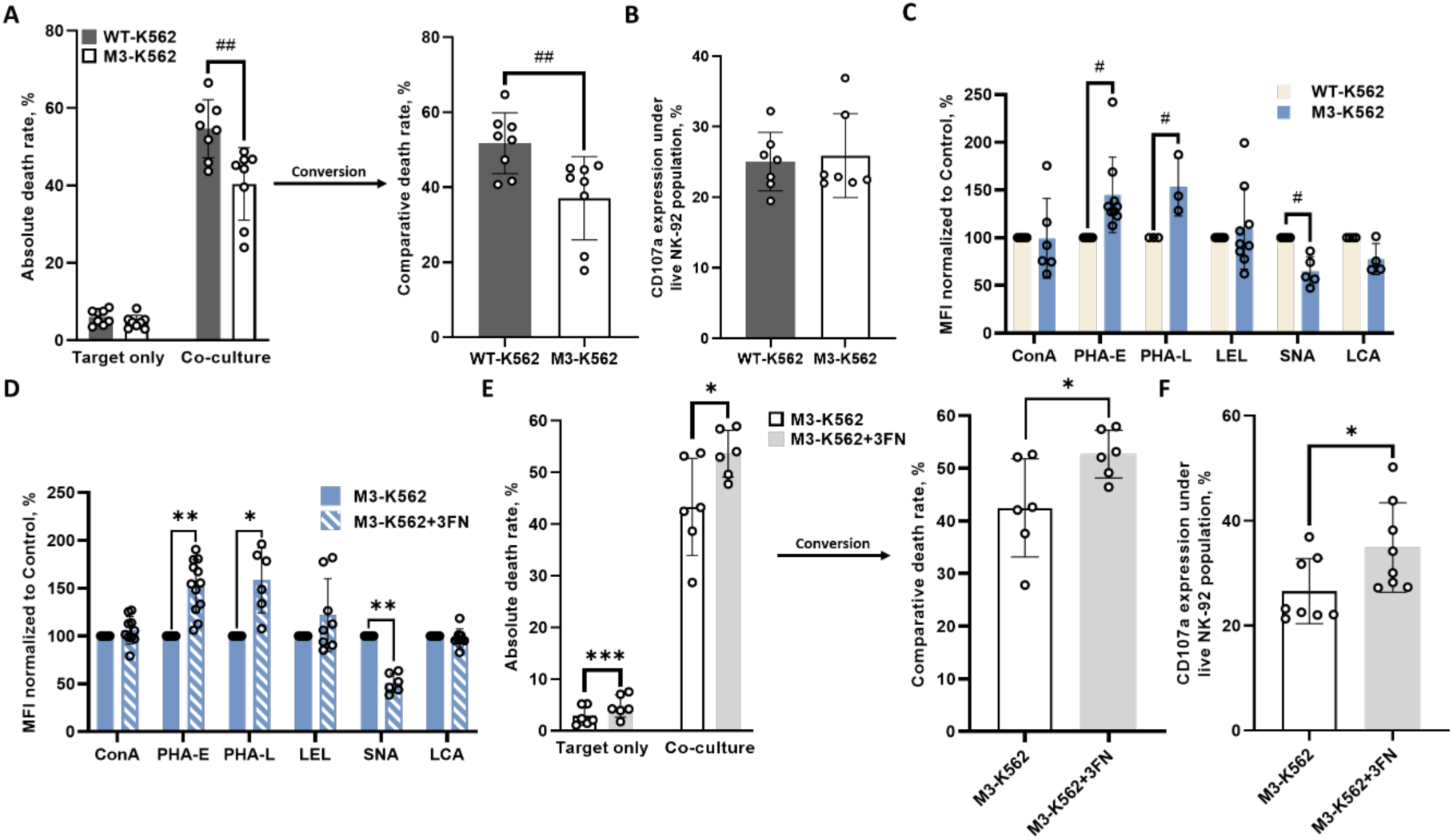
The low NK-susceptibility of M3-K562 cell is not solely determined by its glycomic profile. Comparisons between *MGAT3* overexpressing K562 cells (M3-K562) and its parental cell line (WT-K562) in **(A)** susceptibility to the co-culture with five-fold of NK-92 cells (N = 7), **(B)** the CD107a expression of NK-92 cell during co-culture with K562 cells (N = 7), and **(C)** profiles of cell-surface glycan (N = 3-9). Unpaired Student’s t-test (#) was used to compare the means between WT- and M3-K562 cells. **(D)** Flow cytometry of M3-K562 cells using staining of representative lectins to compare the glycomic profile of cell surface between vehicle control and 3FN-treated samples (N = 7-12). **(E)** Cytotoxicity assays of NK-92 cell co-incubated with M3-K562 pre-treated with 3FN or not (N = 6). **(F)** The CD107a expression of NK-92 cells after the co-culture with K562 cells M3-K562 pre-treated with 3FN or not. Paired Student’s t-test (*) was used to compare the means between vehicle control and 3FN-treated M3-K562 cells.

### The expression of cell surface HLA-G cannot affect the NK-susceptibility of K562 cells

We further studied if the expression of immunomodulator HLA-G on K562 cell surface can affect its NK-susceptibility. K562 cells were transfected to express either wildtype (K562-HLA-G^wildtype^) or mutant (N110Q) HLA-G (K562-HLA-G^N110Q^) using electroporation (Fig. S2A). Under denaturing conditions, only full-length HLA-G isoform HLA-G1 (38 kDa) was detected by the clone 4H84 in the protein extract of first trimester chorionic villi, while another HLA-G isofroms were observed in both K562-HLA-G^wildtype^ and K562-HLA-G^N110Q^ (Fig. 3A). For K562-HLA-G^wildtype^, in addition to the expected translation of the HLA-G1 (38 kDa, black triangle), the α1 subunit of HLA-G1 (∼20 kDa, red triangle) was also blotted (Fig. 3A). Similarly, two proteins with lower molecular weight at ∼36 kDa (white triangle) and ∼18 kDa (orange triangle) were expressed by K562-HLA-G^N110Q^, indicating the absence of *N*-linked glycan (Fig. 3A). Flow cytometry with both cell surface and intracellular staining and immunoprecipitation further confirmed that HLA-G1 can be expressed on the cell surface by K562 cell in correct conformation becasue it can be immunoreacted with clone MEM-G/9 (Fig. 3B) and immunoprecipitated by clone G233 (Fig. S2E). The flow cytometric analyses of K562 transfectants using different clones of anti-HLA-G antibody (Fig. S2B) showed that the proportion of MEM-G/9 positive population of K562-HLA-G^wildtype^ was much higher than that of K562-HLA-G^N110Q^; however, a certain proportion of 4H84 positive population can be observed in both K562 transfectants (Fig. 3B). Statistical analysis further confirmed that the the proportion of MEM-G/9 positive population (indicating membrane-bound HLA-G1) of K562-HLA-G^wildtype^ was significantly higher than that of K562-HLA-G^N110Q^, while there was no significant difference in the proportion of 4H84 positive population (indicating any unfolded HLA-G fragments containing α1 subunit) between these two K562 transfectants (Fig. 3C).

**Figure 3.**
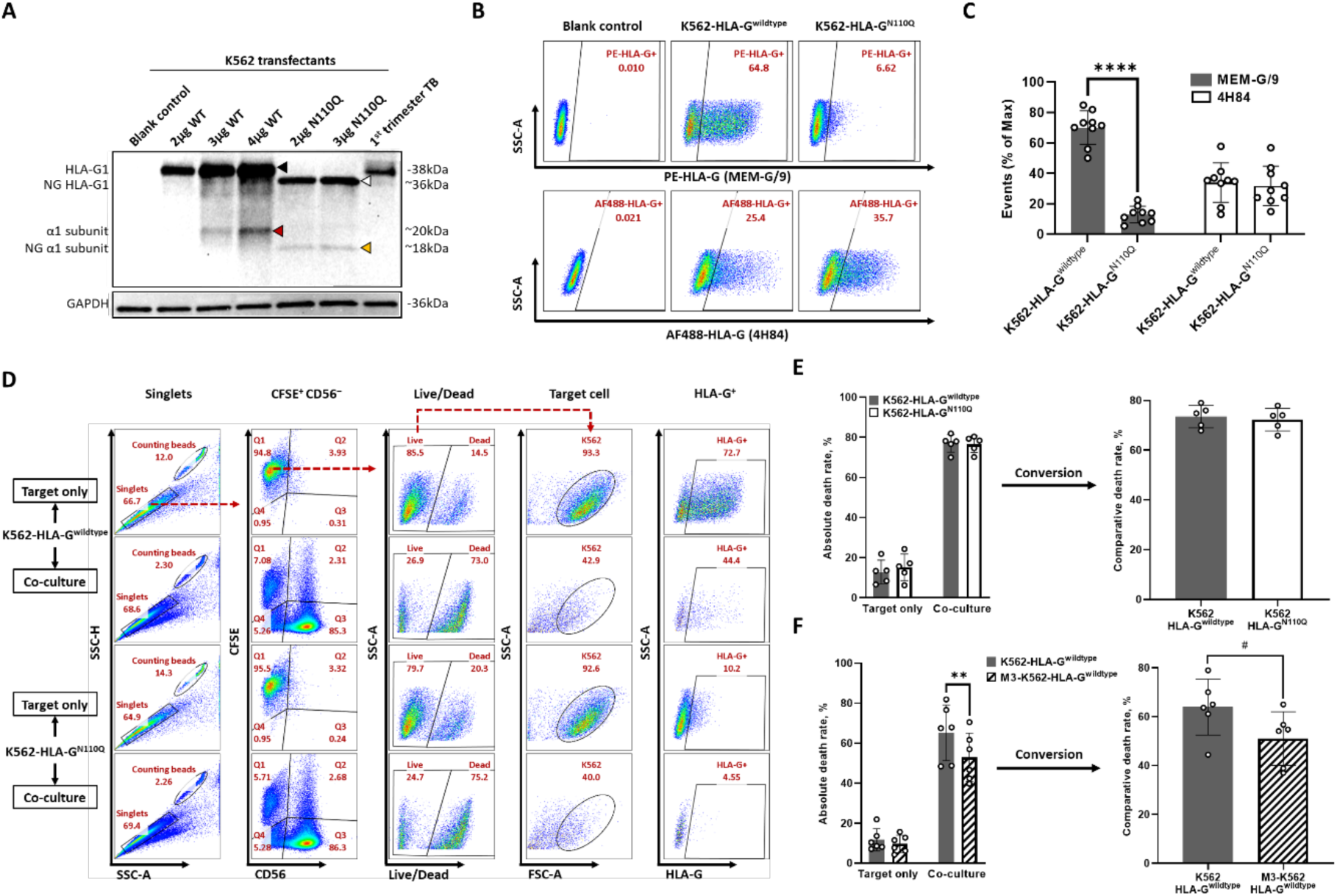
The cell surface expression of HLA-G cannot change the NK-susceptibility of K562 cell. **(A)** Whole cell lysate of K562 transfectants and protein extract of first trimester chorionic villi, were resolved by reducing SDS-polyacrylamide gel electrophoresis and then blotted with clone 4H84. **(B)** Detection of HLA-G isoforms on the cell surface of or inside K562 transfectant by flow cytometric analysis. **(C)** Full length HLA-G1 is abundantly expressed on K562 cell surface after transfection, while unfolded α1 subunits are at the same time accumulating intracellularly. **(D)** Gating strategy for the determination of absolute death rate of K562 transfectants after the co-culture with NK-92 cells. After the exclusion of doublets and debris, both live and dead K562 populations were gated out under CFSE^+^ CD56^−^ population. Under the live population, K562 population was gated in using FSC-A-SSS-A plot and then the HLA-G^+^ population was plotted. The gate for K562 population in FSC-A-SSC-A plot and the baseline of HLA-G^+^ gating area were determined by the “Target only” arm of K562-HLA-G^N110Q^. **(E)** The absolute and comparative death rates of different K562 transfectants after co-culture with NK-92 cells. N = 5 biological replicates. Paired Student’s t-test (*) was used to compare the means between two K562 transfectants. **(F)** The absolute and comparative death rates of K562-HLA-G^wildtype^ and M3-K562-HLA-G^wildtype^ after co-culture with NK-92 cells. N = 6 biological replicates. Unpaired Student’s t-test (#) was used to compare the means between K562-HLA-G^wildtype^ and M3-K562-HLA-G^wildtype^.

Only K562-HLA-G^wildtype^ and K562-HLA-G^N110Q^ were subjected to the co-culture with NK-92 cells to verify if the expression of membrane-bound HLA-G affects the NK-susceptibility of K562 cell, because the introduction of plasmids inherently brought cellular stress to K562 cells, as indicated by the absolute death rates of both transfectants were 4 to 6 fold higher than that of blank control, but no significant difference in absolute death rate between these two K562 transfectants (Fig. S2C, D). No significant difference in either simultaneous or experimental death rate between K562-HLA-G^wildtype^ and K562-HLA-G^N110Q^ was observed, neither was their corresponding comparative death rate, suggesting that these two transfectants exhibited comparable NK-susceptibility (Fig. 3E). Furthermore, M3-K562 transfected to express HLA-G on cell surface (M3-K562-HLA-G^wildtype^), together with K562-HLA-G^wildtype^, were also subjected to flow cytometry-based cytotoxicity assays, and it showed that M3-K562-HLA-G^wildtype^ was still more resistant to the NK-92-mediated cytolysis when compared to K562-HLA-G^wildtype^ (Fig. 3F). A decreased level of ILT2 on NK-92 cell surface after the co-culture with K562-HLA-G^wildtype^ was observed when compared to that co-culture with K562-HLA-G^N110Q^ (Fig. S3A, B), suggesting the binding of NK-92-derived ILT2 to those HLA-G abundantly expressed on K562-HLA-G^wildtype^ cell surface during the cell-cell interaction.

## DISCUSSION

Although much is known about receptor-ligand interactions that govern NK cell-mediated target cell lysis [44], other pathways that influence cancer cell sensitivity or resistance to NK cells have not been fully understood. Aberrant glycosylation is a defining feature of malignancy, allowing tumor cells to evade NK cell surveillance through altered cell surface glycan structures [45,46]. Previous study has provided supporting evidence that differernt types of *N*-linked glycans exhibit distinct impacts on plasma membrane architecture, which further determines diverse cellular activities including growth, maintenance, and stress signaling processes [47]. Immune cells express a wide range of glycan binding receptors capable of detecting and responding to changes in glycan patterns within their environment, consequently, tumor-associated glycan signatures may represent a novel type of immune checkpoint [46,48,49]. This study aimed to determine whether the glycomic profile of target cells influences their susceptibility to cytolysis mediated by NK cell. Our findings demonstrate that changes in *N*-linked glycan processing modulate the NK-susceptibiltiy of K562 cells in a structure-dependent manner. Specifically, the loss of terminal sialylation enhanced K562 cell susceptibility towards NK-92 cell killing. These results provide insight into how alterations in glycosylation of tumor cells may affect their NK-susceptibility and reveal potential immune evasion strategies exploited by malignant cells, highlighting possible targets for cancer immunotherapy.

Our findings all point to a strong overall conclusion that the dominant determinant of NK-susceptibility of K562 cell appears to be the presence of terminal sialic acids rather than simply *N*-linked glycan branching architecture. Only K562 cells treated with either KIF or 3FN became more sensitive to NK-92 cell killing accompanied with an increased CD107a expression by the latter (Fig. 1B, D), and the only common glycomic alteration between these two populations was an decreased level of α2,6-sialylation (Table 1). Morover, the 3FN treatement reversed the high resistance of M3-K562 cell against NK-92 cell killing in despite of elevated levels of GlcNAc branching (Fig. 2D-F), suggesting that terminal sialylation exerts a stronger immunoregulatory effect than glycan branching in this context, or the NK-protective phenotype of bisecting β1,4-GlcNAc observed in the comparison between M3-K562 cells and its parental counterparts (Fig. 2A-C) depends on the surrounding terminal glycan environment. It is known that tumor-associated hypersialylation is relevant in their escape mechanism from the host immune system and is now recognized as a major glyco-immune checkpoint [50]. Several studies have highlighted the role of sialylation in determining the the NK-susceptibility of K562 cell from the perspective of the interaction between glycosylated ligands and inhibitory receptors. Sialic acid polymer that can be incorporated into the plasma membranes to increase cell surface level of Sia, resulting in a status of hypersialylation, has been developed and applied to human cell [51,52]. NK-92 cell was reported to be restrictly activated when co-cultured with K562 cell coated with sialic acid polymers, indicated by an increased recruitment of Src homology 2 domain-containing protein tyrosine phosphatase 1 and a prolonged phosphorylation of Siglec-7 [42]. Leukosialin (CD43 or sialophorin, a negatively charged type I glycoprotein abundantly decorated with *O*-linked glycans) has been identified as the counter-receptor for Siglec-7, of which overexpression on K562 cell can also increase its resistance against NK cell killling in a Siglec-7-dependent manner [53–55]. By contrast, the desialylation of K562 cell by neuraminidase resulted in a significant increase in cytotoxicity mediated by peripheral blood (PB)-NK cell derived from healthy donor; moreover, the blockade of either Siglec-7 or -9 enhances PB-NK-cell mediated cytotoxicity against K562 cell [43], again highlighting the expression of sialylated ligands for both Siglec-7 and -9 on K562 cell surface is playing a key role in modulating its NK-susceptibiltiy of K562. Also using neuraminidase, Daly et al found that expanded PB-NK cells co-cultured with desialylated K562 cell show elevated levels of CD107a, tumor necrosis factor-α, and interferon-γ, suggesting that the desialylation of malignant cell is a promising approach to enhance NK cell-mediated tumor responses [18]. Unlike KIF, even though inducing marked remodelling of cell-surface glycan (Fig. 1A), SWA did not completely abolish complex glycan maturation nor strongly reduce terminal sialylation on K562 cell surface, and the NK-susceptibility of SWA-treated K562 cells remained comparable as the control (Table 1), again higlighting the pivotal role of terminal sialylation in determining the NK-susceptibility of K562 cell. Greco et al have demonstrated that *N*-linked glycans can protect tumors from chimeric antigen receptor-engineered T cell killing by interfering with proper formation of immunological synapse and reducing transcriptional activation, while inhibiting the *N*-linked glycans synthesis in pancreatic adenocarcinoma enhanced chimeric antigen receptor-engineered T cells activity in different xenograft mouse models of pancreatic adenocarcinoma [56]. Simillarly, NK cells can also form immunological synapses to recognize transformed cancer cells based on cell-cell interaction, and secret granzymes and perforin across the synapse to facilitate the cytolysis of cancer cells [57–59]. Future study on whether KIF- or 3FN-treated K562 cells promote the formation of immunological synapse in a similar mechanism would be helpful in understanding the essence of their altered NK-susceptibility.

**Table 1.** Glycomic and phenotypic alteration of JEG-3 cell after inhibitor treatment.

| Compared to control | Lectin binding to K562 cell surface |  |  |  |  |  | NK phenotypes |  |
| --- | --- | --- | --- | --- | --- | --- | --- | --- |
|  | *ConA* | *PHA-E* | *PHA-L* | *LEL* | *SNA* | *LCA* | *NK-susceptibility* | *CD107a* expression by NK-92 |
| **Inhibition of de-mannosylation** |  |  |  |  |  |  |  |  |
| **KIF** | ↑ | ↓ | ↓ | N.S. | ↓ | ↓ | ↑ | ↑ |
| **SWA** |  |  |  |  | N.S. | ↑ | N.S. | ↑ |
| **Pan-inhibition of fucosylation** |  |  |  |  |  |  |  |  |
| **2FF** | N.S. | ↑ | N.S. | N.S. | N.S. | ↓ | N.S. | N.S. |
| **Pan-inhibition of sialylation** |  |  |  |  |  |  |  |  |
| **3FN** | N.S. | ↑ | ↑ | N.S. | ↓ | N.S. | ↑ | ↑ |

**Note:** Arrows denoted increased (↑), or decreased (↓) expression or abundance compared to the control. N = 6-12. N.S., non-significant.

We also studied whether the expression of immunomodulator HLA-G affects the NK-susceptibility of K562 cell. For cytotoxicity assays, it was the first study using K562 cell transfected with non-glycosylated HLA-G as the control because we took stresses brought to K562 cell from both electroporation and expression of exogenous protein (Fig. S2D) into consideration. Although we observed that the expression of cell surface HLA-G1 alone is insufficient to suppress NK-92 cell cytotoxicity in K562 cells (Fig. 3E), probably because NK-92 cells are relatively insensitive to the classical inhibitory HLA signaling which can be observed in decidual NK cell [60], or the lacking of another inhibitory receptor for HLA-G KIR (killer immunoglobulin receptor) which has been demonstrated to play a role in educating both decidual and resting human NK cell [61,62]. At the same time, M3-K562 cell maintained it high resistance against NK-92 cell killing when undertaking the stress from expression of HLA-G when compared to wildtype K562 (Fig. 3F), highlighting the profound influence of remodeling of cell-surface glycans on the NK-susceptibility of K562 cell. However, Salzberger et al have demonstrated that the glycosyaltion inhibition of HLA molecule impair its binding with inhibitory receptor KIR expressed on NK cell surface, and latest evidence further confirmed that defective HLA-G1 glycosylation can disrupt the NK cell tolerance mediated by Siglec-7 at the maternal-fetal interface [63,64]. Therefore, further study is required to answer if the immunoregulatory function of HLA-G is inhibited when presented within an glycoenvironment modified by the overexpression of gene *MGAT3*, rather than intrinsically suppressive by itself.

Overall, this is the first study to clearly demonstrate that, the NK-susceptibility of K562 cells is determined dominantly by the presence of terminal sialic acids rather than the GlcNAc branching architecture in *N*-linked glycan or the accumulation of oligomannose type *N*-linked glycan. Therefore, glycoengineering strategies targeting sialylation may overcome otherwise protective glycan architectures expressed on tumor cell surface.

## AUTHOR CONTRIBUTIONS

Zhengyuan Huang: responsible for experimental design, data acquisition, data analysis, and manuscript writing; Qiqi Li: responsible for experimental design, data analysis, graphing, and manuscript writing; Alex. T. H. Cocker: participated in the writing of the manuscript and performed data analysis; Hugh Brady: responsible for revision of the manuscript; Mark R. Johnson: conceptualised the research, project leaders, and participated in critical revision of the manuscript.

## ACKNOWLEDGEMENTS

We would like to express gratitude to Borne (registered charity number 1167073) and the Overseas Study Program of Guangzhou Elite Project (approval number S. J. 2018, No. 3).

## CONFLICT OF INTEREST STATEMENT

The authors declare no conflicts of interest.

---

**Figure S1.**
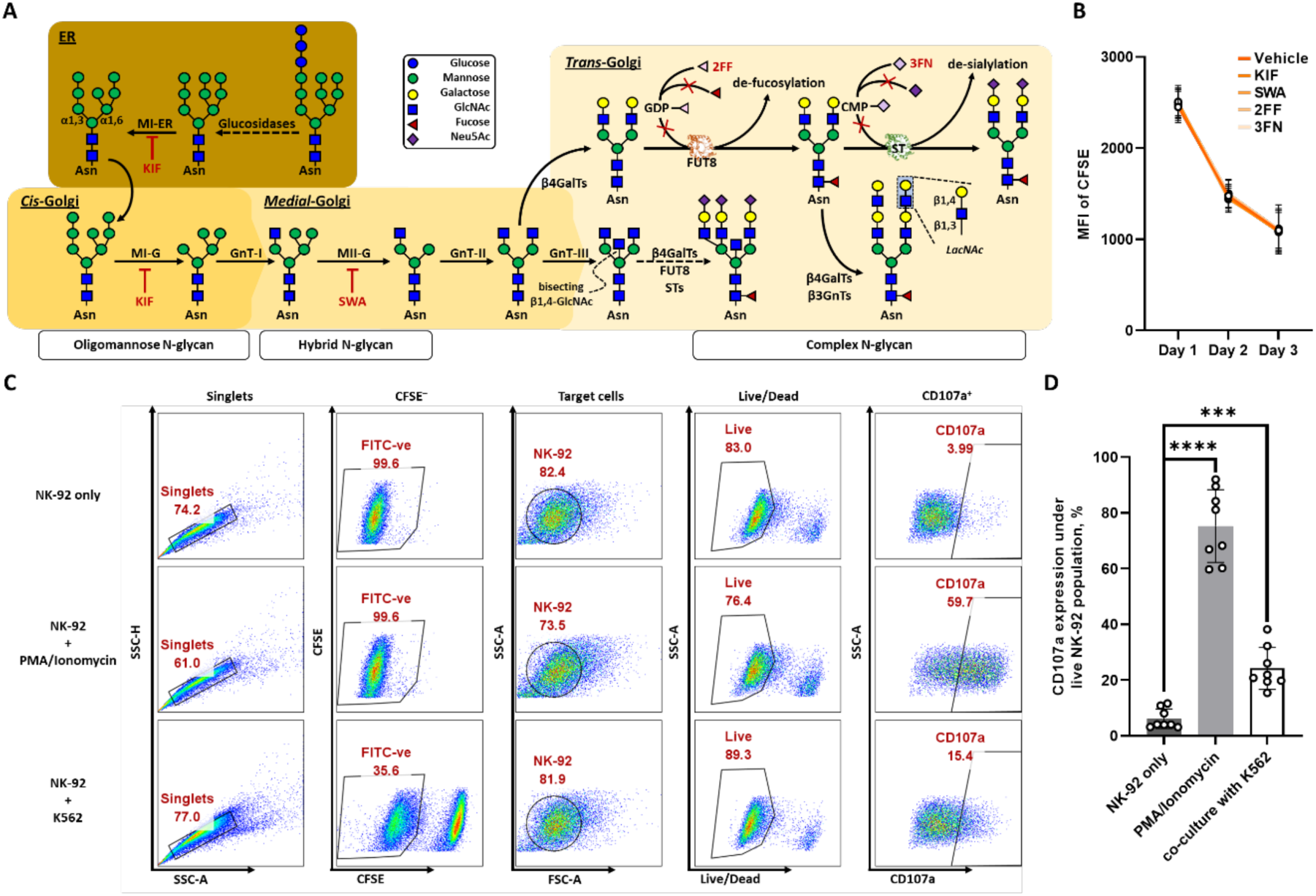
**(A)** Simplified N-glycosylation pathway in mammalian cell with highlighted sites of KIF, SWA, 2FF, and 3FN action and the representative structures expected. β3GnT, β-1,3-N-acetylglucosaminyltransferase; β4GalT, β-1,4-galactosyltransferase; CMP, cytidine monophosphate; FT, fucosyltransferase; FUT8, α-1,6-fucosyltransferase; GDP, guanosine diphosphate; GnT, N-acetylglucosaminyltransferase; MI-ER/-G, α-mannosidase I resident in ER/Golgi apparatus; MII-G, α-mannosidase II resident in Golgi apparatus; Neu5Ac, N-acetylneuraminic acid; ST, sialyltransferase. **(B)** Flow cytometry of vehicle control and inhibitor-treated K562 cells to determine the CFSE fading over three-day incubation to compare the rate of cell proliferation. One-way ANOVA with Bonferroni’s post-test was used to compare the MFI means of CFSE among samples at each timepoint. **(C)** Gating strategy for the determination of CD107a expression by NK-92 cells with PMA/ionomycin stimulation or co-incubated with K562 cells. CFSE^−^ populations were first plotted under singlets, and debris were excluded in FSC-A-SSC-A plot. Dead populations were excluded so that the levels of CD107a^+^ population were determined only under live NK-92 singlets. The gate for NK-92 population in FSC-A-SSC-A plot and the baseline of CD107a positive gating area were based on the “NK-92 only” arm. **(D)** The CD107a expression of NK-92 cell under different conditions compared to blank control. N = 8 biological replicates. Paired Student’s t-test (*) was used to compare the means between blank control and sample with stimulation.

**Figure S2.**
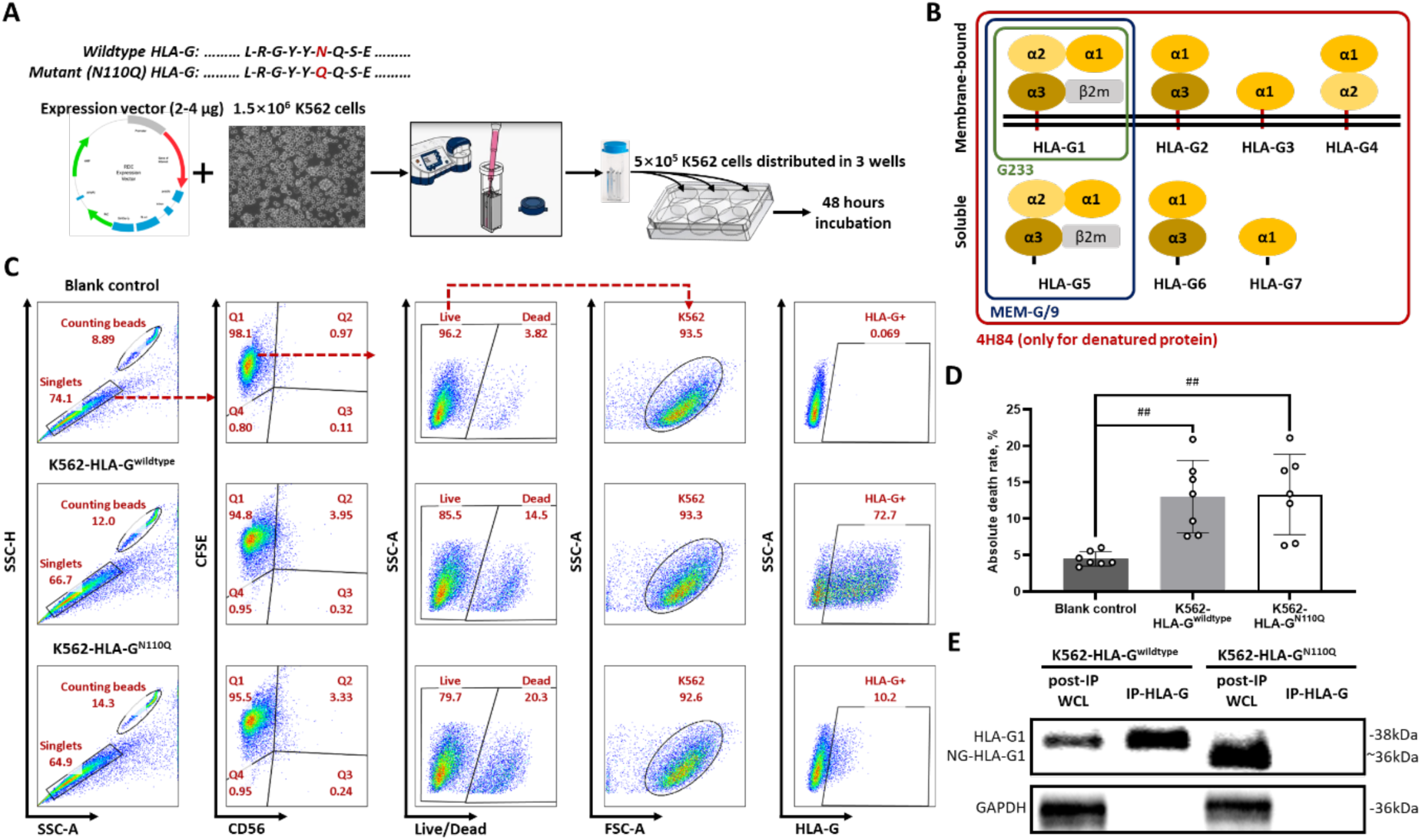
**(A)** Plasmid transfection in K562 cell using electroporation. K562 cells in logarithmic growth were collected, counted, resuspended in electroporation buffer, and combined with gene expression vectors. The mixture was transferred to a cuvette and electroporated on the Nucleofector® 2b. Afterwards, pre-warmed R-10 media was added, and the cell suspension was equally transferred to a 6-well plate and incubated for 48 hours. **(B)** Schematic representation of the various HLA-G isoforms and binding preferences of three different anti-HLA-G mAbs. The primary transcript of the HLA-G gene is alternatively spliced, resulting in at least four membrane-bound and three secreted isoforms. HLA-G isoforms that are framed indicate that they are preferentially recognized by corresponding anti-HLA-G mAbs [1,2]. Especially, clone 4H84 can only be immunoreacted with denatured HLA-G molecules. **(C)** Gating strategy for determining both absolute death rate and HLA-G expression of K562 transfectants. After the exclusion of doublets, live and dead K562 populations were gated out under CFSE^+^ population, respectively. Under the live K562 cell population, debris was excluded using FSC-A-SSC-A plot and then HLA-G positive population were plotted out. The gate for K562 population in FSC-A-SSC-A plot and the baseline of HLA-G positive gating area were based on the “Blank control” arm. **(D)** The stress brought by transfection impairs K562 vitality. N = 7 biological replicates. Data are expressed as the mean and SD. One-way ANOVA (#) with Bonferroni’s post-test was used to compare the means of absolute death rate. **(E)** The immunoprecipitation of HLA-G1 from whole cell lysate of K562 transfectant followed by Western blotting of HLA-G isoforms. HLA-G1 was first immunoprecipitated by clone G233 from total cell lysates of either K562-HLA-G^wildtype^ or K562-HLA-G^N110Q^, and then immunoprecipitated HLA-G1 (IP-HLA-G1) and whole cell lysate after immunoprecipitation (post-IP WCL) were resolved by reducing SDS-polyacrylamide gel electrophoresis and then blotted with clone 4H84. The Western blot of GAPDH was used to indicate the specificity of clone G233 to native HLA-G1. Representative blots are shown.

**Figure S3.**
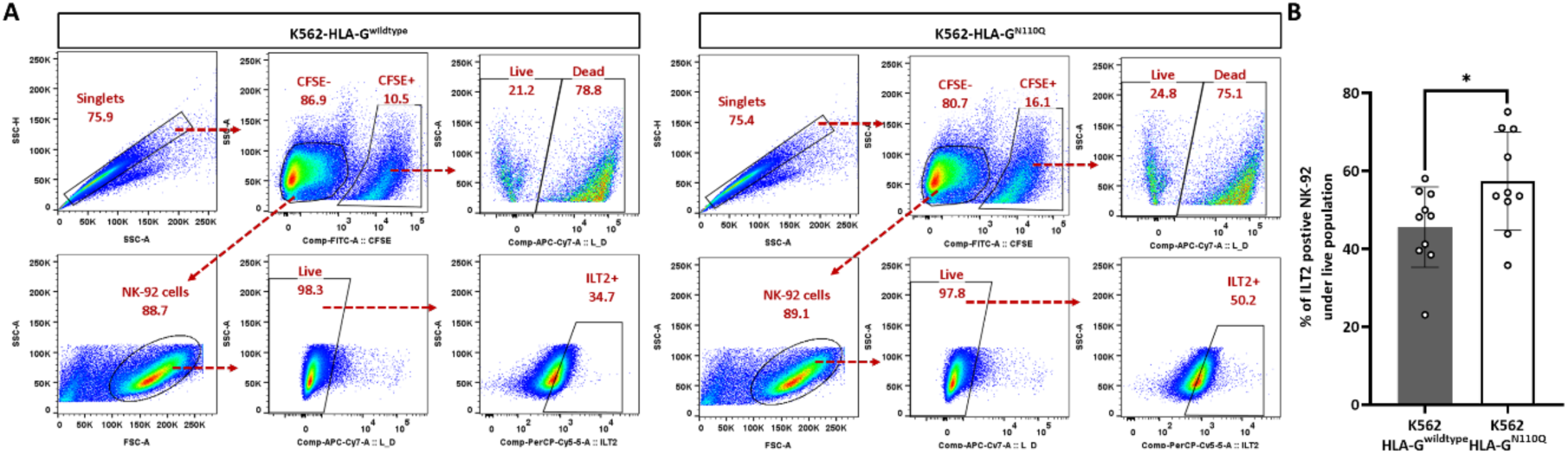
**(A)** Gating strategy determining the proportion of ILT2 positive NK-92 cells after co-culture with K562 transfectants. After the exclusion of doublets, CFSE^−^ population was plotted as NK-92 cells. Debris were excluded using FSC-A-SSC-A plot under the CFSE^−^ population, and then live NK-92 singlets were plotted. Eventually, ILT2^+^ population was plotted under live NK-92 singlets. Live and dead K562 populations were plotted under the CFSE^+^ population, respectively. The gate of ILT2^+^ population was determined by the “K562 only” arm. Representative plots are shown. **(B)** Representative data for percentage of ILT2 positive NK-92 under live population. N = 10 biological replicates. Unpaired Student’s t-test (#) was used to compare the means of percentages of ILT2 positive population under live NK-92 population.

## Notes

### Competing Interest Statement

The authors have declared no competing interest.

### Funding

- Borne Foundation
- Chelsea and Westminster Hospital NHS Foundation Trust

