## Supplementary Tables for "Cell-surface *N*-linked glycans manipulation of K562 cells for augmented susceptibility to natural killer cell killing"

**Table S1. Representative plant lectins for cell surface glycan analysis.**

| **Lectin** | **Glycan-binding specificity** | **References** |
| --- | --- | --- |
| *Canavalia ensiformis* agglutinin (ConA) | Mannose, glucose | [1] |
| *Lens culinaris* lectin (LCA) | Core α-1,6-linked fucose |  |
| *Lycopersicon esculentum* lectin (LEL) | Polylactosamine chains |  |
| *Phaseolus vulgaris* erythronagglutinin (PHA-E) | Bisecting β-1,4-GlcNAc branch |  |
| *Phaseolus vulgaris* leukoaggulutinin (PHA-L) | β-1,6-GlcNAc branch |  |
| *Sambucus nigra* agglutinin (SNA) | α-2,6-linked sialic acid |  |

**Table S2. Inhibitors for different N-glycan processing enzymes.**

| **Inhibitor** | **Structure** | **Targeting enzyme** | **Mechanism** | **Working conc.** |
| --- | --- | --- | --- | --- |
| Kifunensine (KIF) | 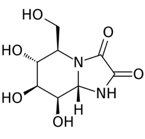 | α-mannosidase I resident in both ER and Golgi (MI-EG) | Inhibition of α-mannosidase I resident in ER and Golgi | 10 μg/mL |
| Swainsonine (SWA) | 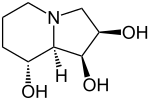 | α-mannosidase II resident in Golgi apparatus (MII-G) | Inhibition of Golgi α-mannosidase II | 10 μg/mL |
| 2F-Peracetyl-Fucose (2FF) | 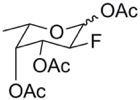 | Fucosyltransferases (FTs) | Metabolic transformation into a GDP-fucose mimetic | 500 µM |
| 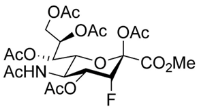3F_AX_-Peracetyl Neu5Ac (3FN) |  | Sialyltransferases (STs) | Inhibit ST inhibitor in a donor substrate CMP-Neu5Ac-competitive manner | 300 µM |
